# Poorly differentiated XX/XY sex chromosomes are widely shared across skink radiation

**DOI:** 10.1101/2020.08.29.273524

**Authors:** Alexander Kostmann, Lukáš Kratochvíl, Michail Rovatsos

## Abstract

Differentiated sex chromosomes are believed to be evolutionarily stable, and their emergence was suggested to lead to a remarkable increase in the diversification rate and in disparity in such groups as birds, mammals and snakes. On the other hand, poorly differentiated sex chromosomes are considered to be prone to turnovers. With around 1.700 currently known species forming *c*. 15% of reptile species diversity, skinks (family Scincidae) are a very diverse group of squamates known for their large ecological and morphological variability. Skinks generally have poorly differentiated and cytogenetically hardly distinguishable sex chromosomes and their sex determination was suggested to be highly variable. Here, we determined X-linked genes in the common sandfish (*Scincus scincus*) and demonstrate that skinks have shared the same homologous XX/XY sex chromosomes across their wide phylogenetic spectrum for at least 85 million years, approaching the age of the highly differentiated ZZ/ZW sex chromosomes of birds and advanced snakes. Skinks thus demonstrate that even poorly differentiated sex chromosomes can be evolutionarily stable and that large diversity can emerge even in groups with poorly differentiated sex chromosomes. The conservation of sex chromosomes across skinks allows us to introduce the first molecular sexing method widely applicable in this group.

## Introduction

Organisms do not share a common mechanism for sex determination. Under genotypic sex determination (GSD), the sex of an individual is set at conception by its sex-specific genotype, i.e. by the sex-specific combination of sex chromosomes. GSD is very common in animals and has evolved in them multiple times, it is estimated that this has occurred independently up to 40-times just within amniotes [1]. Sex chromosomes are thus a notable example of convergent evolution. Surprisingly, in spite of over a century of research [2], the adaptive significance and consequences of sex chromosomes and their differentiation, the progressive cessation of recombination and divergence of sequences between chromosomes in a sex chromosome pair, are still rather controversial. Sex chromosomes primarily determine the sex of an individual and ensure a stable sex ratio at conception; however, the sex-specific specialization and particularly, the role in the resolution of the conflict between sexes over trait expression have been considered crucial for differentiation and evolutionary stability of sex chromosomes [1,3]. Nevertheless, there are more ways to resolve sexual conflict, the most important in vertebrates being the control of the expression of sex-specific traits by sex hormones [4,5]. Species without sex chromosomes such as vertebrates with environmental sex determination (ESD) are considerably sexually dimorphic as well [6], and the importance of sex chromosomes for the resolution of sexual conflict is still highly debated [7].

Next to the above-mentioned adaptive advantages, organisms with differentiated sex chromosomes also incur important costs. Higher mortality and a reduced lifespan in individuals of the heterogametic sex leading to a biased adult sex ratio attributed to degeneration of the Y and W were reported across animal lineages [8,9]. The emergence of differentiated sex chromosomes could also have important macroevolutionary consequences. It was noted that it proceeded in parallel with the remarkable increase in diversification rates within birds, mammals and snakes [10]. The superradiation of the clades with differentiation of sex chromosomes was suggested to be attributable to the so-called ‘fast Z’, and likewise ‘fast X’ phenomena, i.e. the increased evolutionary rate of the Z-specific and likewise X-specific genes relative to the autosomal or pseudoautosomal genes, potentially enabling higher diversification in lineages with differentiated sex chromosomes.

The causes of the unequal rate of differentiation of sex chromosomes among lineages are still unresolved. The different selection pressures in males compared to females, led to the traditional theoretical predictions that sex chromosomes should differentiate faster under male heterogamety (XX/XY) than under female heterogamety (ZZ/ZW). The expectation of the faster differentiation (or degeneration) of Y was based on assumptions of a stronger selection in males, male mutation bias, or a smaller effective population size of the Y chromosome with a decreasing male to female ratio in a population [11–13]. However, the very recent model stressing differences in the recombination rates between sexes and among lineages suggested just the opposite pattern: inversions reducing or arresting recombination should be fixed more frequently on Z and W than X and Y chromosomes, and it is therefore expected that the W chromosome degenerates faster than the Y [14]. We clearly need more empirical data on the rate of differentiation of sex chromosomes in different lineages with the opposite heterogamety to solve this controversy.

Among vertebrates, sex chromosomes were traditionally believed to be stable in endotherms, i.e. in mammals and birds, but unstable in fish, amphibians and non-avian reptiles [15]. For reptiles, multiple transitions between ESD and GSD in both directions were expected [16]. This traditional paradigm was shaken only recently by a series of molecular studies documenting that many reptile lineages, specifically advanced snakes, lacertid lizards, anguimorphan reptiles, iguanas and softshell turtles possess in fact highly differentiated and long-term, stable sex chromosomes [17–19]. The minimal age of sex chromosomes in these lineages is approximately between 80 and 180 MY [20,21] and thus comparable to the minimal age of sex chromosomes of birds (100-120 MY) and viviparous mammals (165 MY) [22–24]. For squamates, it was suggested that sex chromosomes evolved a long time ago independently, possibly from the ancestral ESD, in most of their major clades. Only dragon lizards, geckos and possibly skinks were considered exceptional due to extensive variability in sex determination systems [1,25].

With around 1.700 currently known species forming *c*. 15% of reptile species diversity, skinks (family Scincidae) are a very diverse, nearly cosmopolitan group of squamates known for their large ecological and morphological variability [26]. They include terrestrial, even subterraneous, arboreal, and semiaquatic forms with numerous transitions to viviparity, limblessness, and nocturnality and significant variability in body size [27]. In spite of decades of cytogenetic research, sex chromosomes have been determined only in about a dozen skink species across their enormous diversity [28–31]. Variability of sex determination in skinks was suggested for a long time. Previously Donnellan [32] suggested that sex chromosomes are not homologous across skink lineages based on chromosome morphology. However, chromosome morphology is not the decisive criterion of (non)homology of sex determination systems. The situation was complicated further by reports of an environmental influence on sex in certain species of skinks, leading to the conclusion that some members of this lineage have ESD and hence no sex chromosomes [33–35]. Nevertheless, many earlier reports of ESD were found to be unreliable based on recent cytogenetic or molecular evidence [36–39], which caused an overestimation of the number of GSD to ESD transitions among amniotes and undermined the long-term stability of GSD.

We decided to shed light on this long-lasting debate. By comparison of the male and female genomes, we identified X-specific genes, i.e. genes present on X but missing on the Y chromosome, in the common sandfish (*Scincus scincus*), one of the few skink species with cytogenetic evidence for sex chromosomes [40]. After validation of X-specificity by quantitative PCR (qPCR) in this species, we performed a molecular test of the homology of sex chromosomes across skinks and their nearest outgroups.

## Material and Methods

### Studied material

Tissue samples (blood from living individuals and muscle or tissue from the tip of the tail from preserved specimens) were collected from one male and one female individual of 13 representative species of skinks and their close outgroups represented by members of the families Cordylidae, Gerrhosauridae and Xantusiidae, who along with skinks form the clade Scincoidea (Table S1). Selection of species was done according to their phylogenetic position to cover all major skink lineages [41] and availability of high-quality tissue samples of both sexes for DNA isolation. We were not able to include the African subterranean subfamily Acontinae, where sexed material was not available to us. Genomic DNA was isolated from blood or tail tissue using a Qiagen DNeasy Blood and Tissue kit (Qiagen). All experimental procedures were carried out under the supervision and with the approval of the Ethics Committee of the Faculty of Science, Charles University in Prague, followed by the Committee for Animal Welfare of the Ministry of Agriculture of the Czech Republic (permission No. 29555/2006-30).

### *Chromosome preparations and cytogenetic analysis of* Scincus scincus *and* Tropidophorus baconi

Metaphase chromosomes were prepared from whole blood cell cultures according to the protocol described in Mazzoleni et al. [42] in widely phylogenetically distant skinks *Scincus scincus* (Scincinae) and *Tropidophorus baconi* (Lygosominae). The chromosomes were Giemsa-stained and photographed using a Provis AX70 fluorescence microscope equipped with a digital camera (Olympus DP30BW). C-banding was applied following Sumner [43] to uncover heterochromatic regions, which are often accumulated on differentiated sex chromosomes. Karyograms were prepared using Ikaros software (MetaSystems, Germany). To identify sex-specific differences in sequence composition, we performed comparative genomic hybridization (CGH) using a previously described protocol [36]. Also, we examined the distribution of the rDNA loci as the sex chromosomes of *Scincus scincus* were identified by differences in numbers of nucleolar organizing regions (NORs) between sexes [40], using fluorescence *in situ* hybridization (FISH). The probe for the rDNA loci was prepared from a plasmid (pDm r.a51#1) with an 11.5-kb insertion, encoding the 18S and 28S rRNA units of *Drosophila melanogaster* [44]; this protocol is explained in detail in Rovatsos et al. [36].

### Genome coverage analysis in S. scincus

Genomic DNA isolated from blood samples of one male and one female of *Scincus scincus* was sequenced at high coverage by Novogene (Cambridge, UK) in Illumina HiSeq2500 platform, with 150 base pairs (bp) pair-end option (DNA-seq). The raw Illumina reads are deposited in Genbank under the NCBI BioProject PRJNA660179. Adapters and low-quality bases from raw reads were trimmed by Trimmomatic [45] and reads shorter than 50 bp were removed. Trimmed reads were checked in FASTQC [46] and MULTIQC [47].

The X-specific genes have half the copy numbers in the genomes of XY males in comparison to XX females. The differences in the copy numbers of X-specific genes between sexes are expected to be proportional to the differences in coverage of the reads from DNA sequencing in Illumina HiSeq platforms [48,49]. Therefore, these loci are expected to have half read coverage in males compared to females, while autosomal and pseudoautosomal loci should have equal read coverage in both sexes. We independently mapped the trimmed Illumina reads from the male and the female sandfish using Geneious Prime (https://www.geneious.com) to a reference dataset of 237,298 exons of the common wall lizard (*Podarcis muralis*), a lizard with a genome assembled to chromosome level and high quality gene annotation [50]. The average read coverage per gene was calculated in each specimen after filtering all exons with unexpectedly high or low coverage (i.e. 4-fold difference from the average coverage). Subsequently, we calculated the ratio of female to male read coverage for each gene, normalized to the total number of assembled reads per specimen [table S2]. We identified X-specific genes as genes with a male to female coverage ratio between 0.35 and 0.65.

Also, X-specific, single copy genes are hemizygous in the heterogametic sex (i.e. in males in *S. scincus*). Therefore, such loci should not have single nucleotide polymorphisms (SNPs) in our map-to-reference assembly from males. We calculated the presence/absence of SNPs in the assembly of the male sandfish in order to validate the X-specificity of genes uncovered by the comparative coverage analysis [table S2].

### Validation of X-specific loci and test of homology of sex chromosomes across skinks by qPCR

The differences in gene copy numbers of the X-specific genes between sexes can be measured by qPCR applied to genomic DNA [10,18]. We applied this technique to further validate X-specificity of a subset of X-specific genes uncovered from bioinformatic analyses. Subsequently, as a test of homology of sex chromosomes across skinks, we tested whether the X-specific genes in *S. scincus* are also X-specific in other skinks and their close outgroups. For qPCR, we designed specific primers for sandfish X-specific genes using Primer3 software [51] for the amplification of a 120–200 bp fragment of three single-copy autosomal genes (*abarb2, eef1a1, mecom*) and 10 X-specific genes [table S3]. The qPCR was carried out in a LightCycler II 480 (Roche Diagnostics) with all samples run in triplicate. The detailed qPCR protocol and the description of the calculation of the relative gene dose between sexes are available in our previous reports [17,52]. Briefly, the gene dosage of each target gene was calculated from crossing point values and was subsequently normalized to the dose of the single-copy reference gene *eef1a1* or *mecom* from the same DNA sample. The relative gene dosage ratio (*r*) between sexes of a given species for each target gene is expected to be close to 0.5 for X-specific genes, 1.0 for (pseudo)autosomal genes and potentially 2.0 for Z-specific genes.

## Results and Discussion

### Cytogenetic analysis

Caputo et al. [40] reported the sex-associated NOR polymorphism in *S. scincus* with two active NORs in females, while only one in males. We confirmed this observation by FISH with the probe specific for rDNA sequences, demonstrating that the sex chromosomes in this species differ both in the transcription activity (NORs) and the number of rDNA repeats [figure 3a]. Nevertheless, the sex-related polymorphism in rDNA loci is not present in the genera *Eumeces* and *Plestiodon* closely related to *S. scincus* [40], nor in *Tropidophorus baconi* [figure 3b]. Thus, it seems that the polymorphism enabling us to distinguish sex chromosomes cytogenetically is an apomorphy of the sandfish lineage. Surprisingly, sex-specific differences in the genome were not detected by CGH in either *S. scincus* [figures 3c,3d], or in *Tropidophorus baconi* [figures 3e,3f]. The lack of sex-specific signal in CGH indicates that the sex chromosomes (if present in *Tropidophorus baconi*) are homomorphic and poorly differentiated.

**Figure 1:**
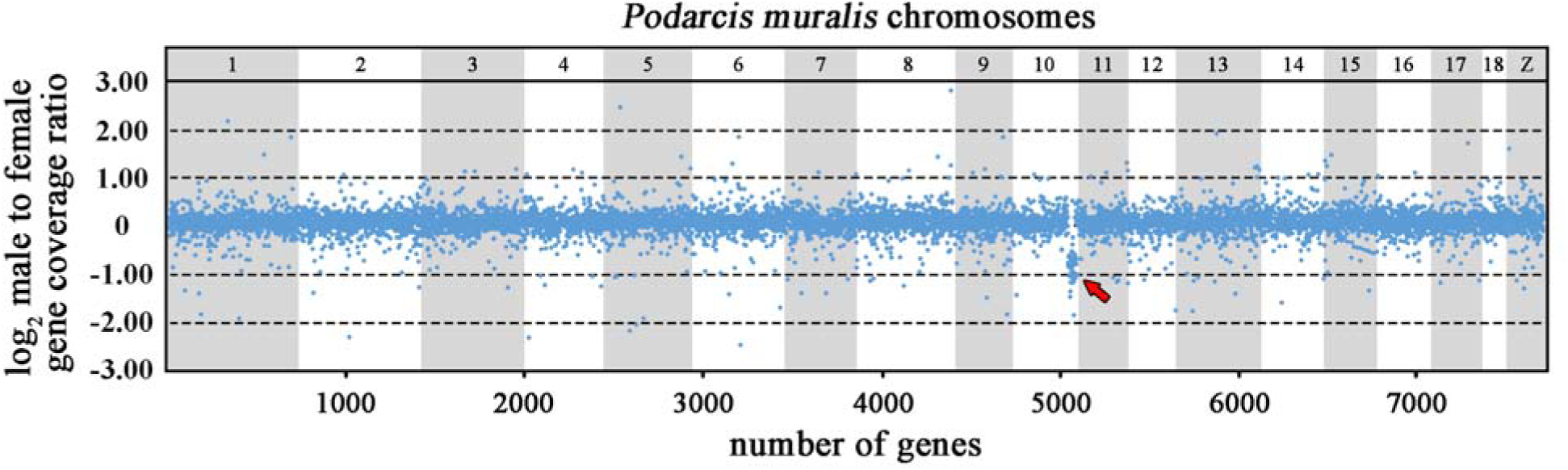
Log_2_-transformed male to female ratios of DNA-seq read coverage per gene in *Scincus scincus*. The genes are illustrated according to the position of their orthologs in the chromosomes of *Podarcis muralis*. Note that the region homologous to PMU10 (indicated by a red arrow) possesses half male to female ratio in read coverage depth, demonstrating that this genome part is X-specific in *Scincus scincus*.

**Figure 2:**
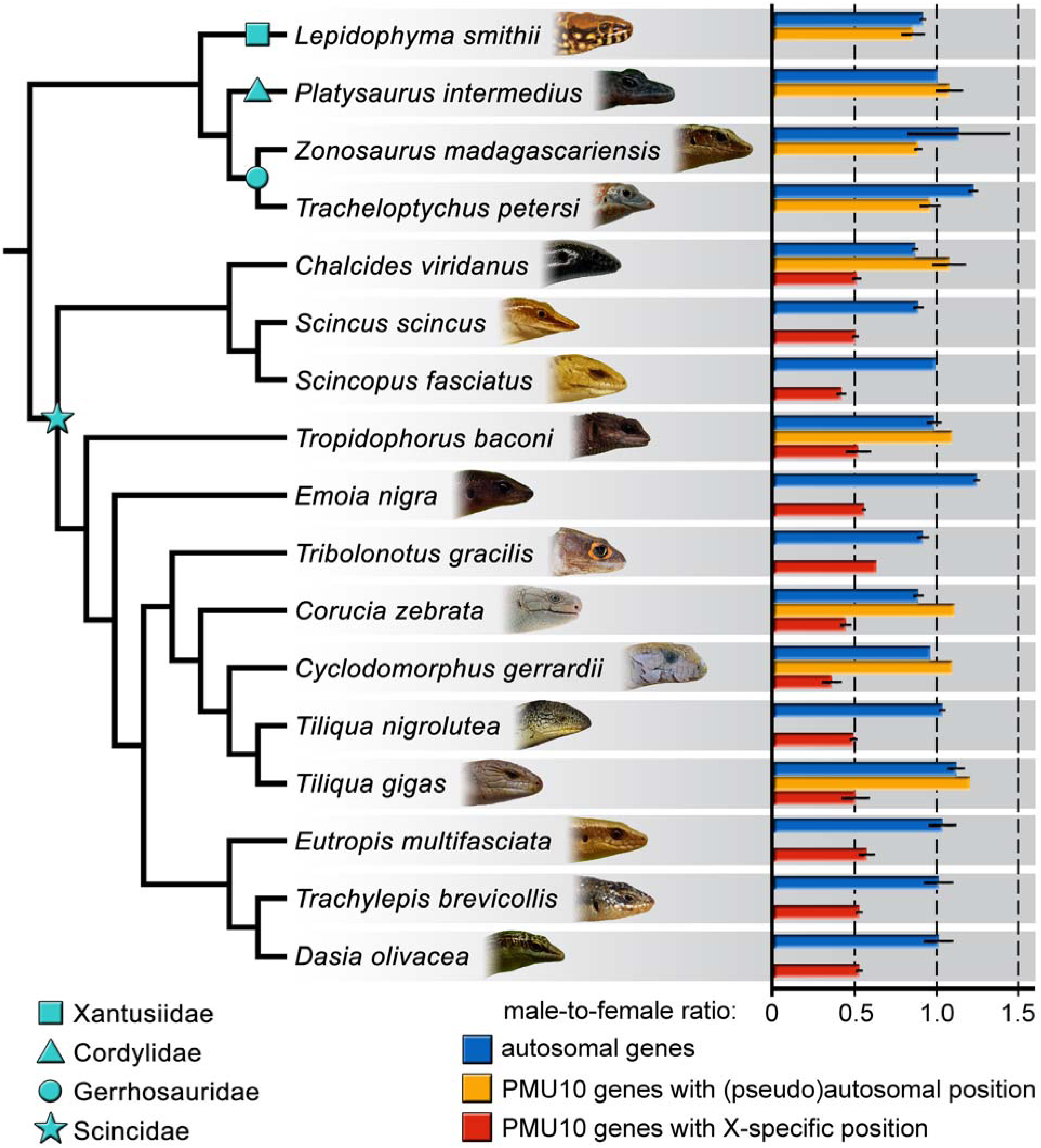
Average relative gene dose ratios between males and females for autosomal genes (blue) and PMU10 genes with (pseudo)autosomal (orange) and X-specific genes (red) across 13 species of skinks (Scincidae) and four outgroup species from the families Cordylidae, Gerrhosauridae and Xantusiidae.

**Figure 3:**
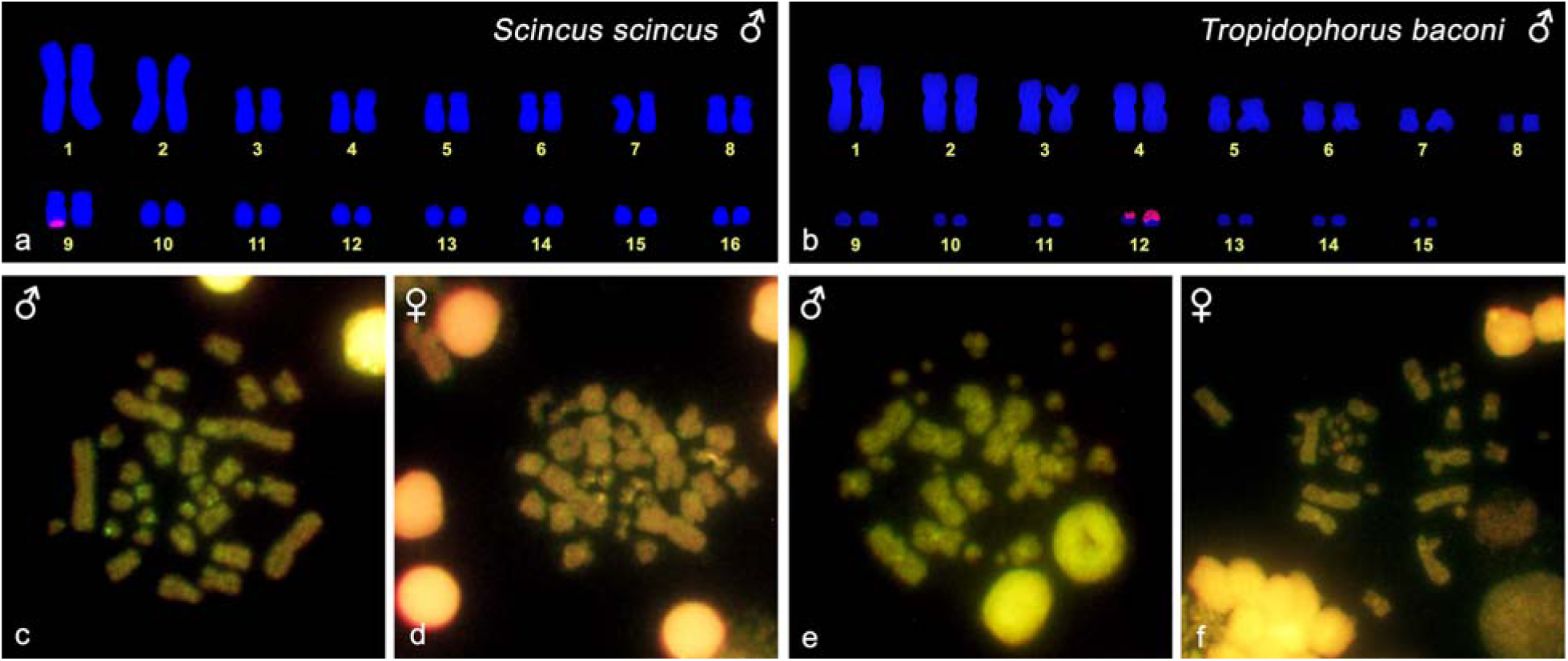
Karyogram reconstruction from metaphase after FISH with the probe for rDNA loci, in male individuals of *Scincus scincus* (a) and *Tropidophorus baconi* (b). Signal of rDNA loci was detected in a single chromosome from the 9^th^ pair in males of *S. scincus*, but in both chromosomes of the 12^th^ pair in *T. baconi*. CGH revealed differences between sexes neither in males, nor in females in both *S. scincus* (c,d) and *T. baconi* (e,f).

### *X-linked genes in* S. scincus

The analysis of exonic genome regions in the common sandfish revealed 560 genes with roughly half coverage in the male in comparison to the female genome (male to female ratio between 0.35-0.65), which is expected for X-specific genes missing on the Y chromosome[figure 1; table S2]. Notably, 169 out of 560 candidate X-specific genes were lacking polymorphisms in males (determined as absence of SNPs in at least 80% of exons in a given gene), which is in accordance with the expected hemizygous state. The homologs of these 169 genes are scattered across all *P. muralis* chromosomes. Notably, 37 of these genes are linked to *P. muralis* chromosome 10 (PMU10), covering a chromosomal region of approximately 7 million base pairs [table S2]. The X-specificity of a subset of 10 genes from PMU10 was validated by comparison of copy number variation between the sexes by quantitative PCR (qPCR) [figure 2; table S3).

The X-linked genes from the region homologous to a part of PMU10 have homologs linked to a region of chicken chromosome 1 (GGA1). As far as is known, the PMU10/GGA1 syntenic block is not involved in sex chromosomes in other amniote lineages [22] supporting the hypothesis that the sex chromosomes in skinks evolved independently from other amniote sex chromosomes. Notably, this chromosomal region contains several genes, which are involved in gonad development or their pathological conditions (*dmc1, ep300, igf1, kitlg, nup107, pdgfb, sbf1, sox10, stra8, sycp3*) and can potentially act as the sex determining gene(s) in skinks. Among these genes, prominent candidates could be the genes *ep300* (E1A binding protein p300), a histone acetyltransferase that regulates transcription via chromatin remodeling, and *sox10* (*sry*-box transcription factor 10), a member of the *sox* (*sry*-related hmg-box) family of transcription factors involved in the regulation of embryonic development. Mouse embryos lacking functional copies of *ep300* exhibit complete XY gonadal sex reversal [53]. In pathological conditions, the ectopic expression of *sox10* in embryonic gonad leads to upregulation of transcriptional targets of the *sox9* gene, triggering the male differentiation pathway, and resulting in sex-reverted XX males in humans and mice [54,55]. A third potential sex determining gene is *sbf1* (SET binding factor 1), a member of the protein-tyrosine phosphatase family involved in cell growth and differentiation which is highly expressed in the brain and testes. Alterations in the *sbf1* sequence or splicing lead to male infertility, impaired spermatogenesis and azoospermia in humans, mice and rats [56–59]. Notably, *sbf1* is nested in the 7 million base pair chromosomal region, enriched with X-specific genes in *S. scincus*.

It was suggested that certain genome sections may tend to be recruited for the function of sex chromosomes, or be added later as a part of neo-sex chromosomes, non-randomly more often than other parts due to the content of genes involved in gonad differentiation (e.g. *amh, dmrt1, sox3*) and sex-specific traits [50–62]. It is true that some syntenic blocks were co-opted for the role of sex chromosomes across amniotes several times, for example the ortholog of GGAZ in birds, in a turtle lineage, and twice independently in geckos, the ortholog of GGA4p which makes up the main part of the sex chromosomes in viviparous mammals, geckos of the genus *Paroedura* and lacertid lizards [22], and GGA28 in anguimorphan lizards and monotremes [18].

However, other lineages such as the caenophidian and non-caenophidian snakes, the xantusiid lizard *Xantusia henshawi*, the pygopodid geckos and skinks [63–66] have evolved sex chromosomes from syntenic blocks not forming sex chromosomes in other amniotes. Nevertheless, in the majority of the studied lineages, only parts of the sex chromosomes have been identified. The accurate identification of sex chromosome gene content and the sex determining genes across all lineages of amniotes that independently evolved sex chromosomes is crucial to conclude whether some syntenic blocks are indeed exceptionally frequently turned into sex chromosomes.

### Poorly differentiated, but highly conserved sex chromosomes across skinks

The copy number variation in the orthologs of X-specific genes of the common sandfish was consistent with their X-specificity in all 13 tested skinks covering a phylogenetic spectrum of 1.665 extant species. Our results support that sex chromosomes are highly conserved across the skink radiation [figure 2], in spite of their poor differentiation. We confirmed the poor stage of differentiation in *Tropidophorus baconi*, where cytogenetic techniques including CGH failed to reveal sex chromosomes, although qPCR confirmed that this species shares XX/XY sex chromosomes with the common sandfish. In support of our results, a recent study in the water skink *Eulamprus heatwolei* revealed an XX/XY sex determination system, with sex chromosomes homologous to the same part of GGA1 [67]. The small X-specific region of skinks seems to be beyond the detection efficiency of molecular cytogenetic methods such as CGH, which explains why sex chromosomes in most skinks were not detected up to now despite several careful cytogenetic studies [68,69].

The orthologs of X-specific genes of the common sandfish showed a pseudoautosomal or autosomal pattern in the outgroups from the three families (Cordylidae, Gerrhosauridae and Xantusiidae) who along with skinks form the clade Scincoidea. ZZ/ZW sex chromosomes containing genes from other regions were reported in a night lizard from the family Xantusiidae [65], while sex determination is not known up to now in any cordylid or gerrhosaurid lizards. The origin of sex chromosomes in skinks can therefore be dated between the split of the monophylum of the three other scincoidean families and skinks around 150 MYA and the divergence of the subfamilies Lygosominae and Scincinae living *c*. 85 millions years ago [70,71].

### Towards resolution of controversies on sex determination in skinks and other reptiles

At least two species of skinks previously reported as ESD were found to have GSD. In *Bassiana duperreyi* the environmental influence on sex was explained by sex reversal [72] and sex-linkage of anonymous molecular markers consistent with XX/XY sex chromosomes were found in *Niveoscincus ocellatus* [73]. Of course, we cannot exclude the possibility that some species of skinks do not share the same sex chromosomes. Rare exceptions from the conserved pattern with likely derived sex chromosomes were found in iguanas, anguimorphan lizards and mammals [18,74–76]. However, the current results show that the sex chromosomes homologous to the sandfish are conserved for a long time across most of the phylogenetic diversity of skinks. Assuming that all species derived from the common ancestor of Lygosominae and Scincinae share the same chromosomes, we can estimate that roughly at least 15% out of nearly 11.000 currently recognized squamate species share sex chromosomes with the sandfish, and that about 60% of squamate species are members of the five lineages with adequate molecular evidence for conservation of sex chromosomes [figure 4]. We assume that GSD will also be frequent in the remaining 40% of squamate species, with sex chromosomes being reported for example in chameleons, agamids and many geckos [22,36,77,78]. Against older predictions, ESD seems to be rather rare representing approximately 5% of species diversity in non-avian reptiles [79]. Future studies should further verify and increase precision of these estimations.

**Figure 4:**
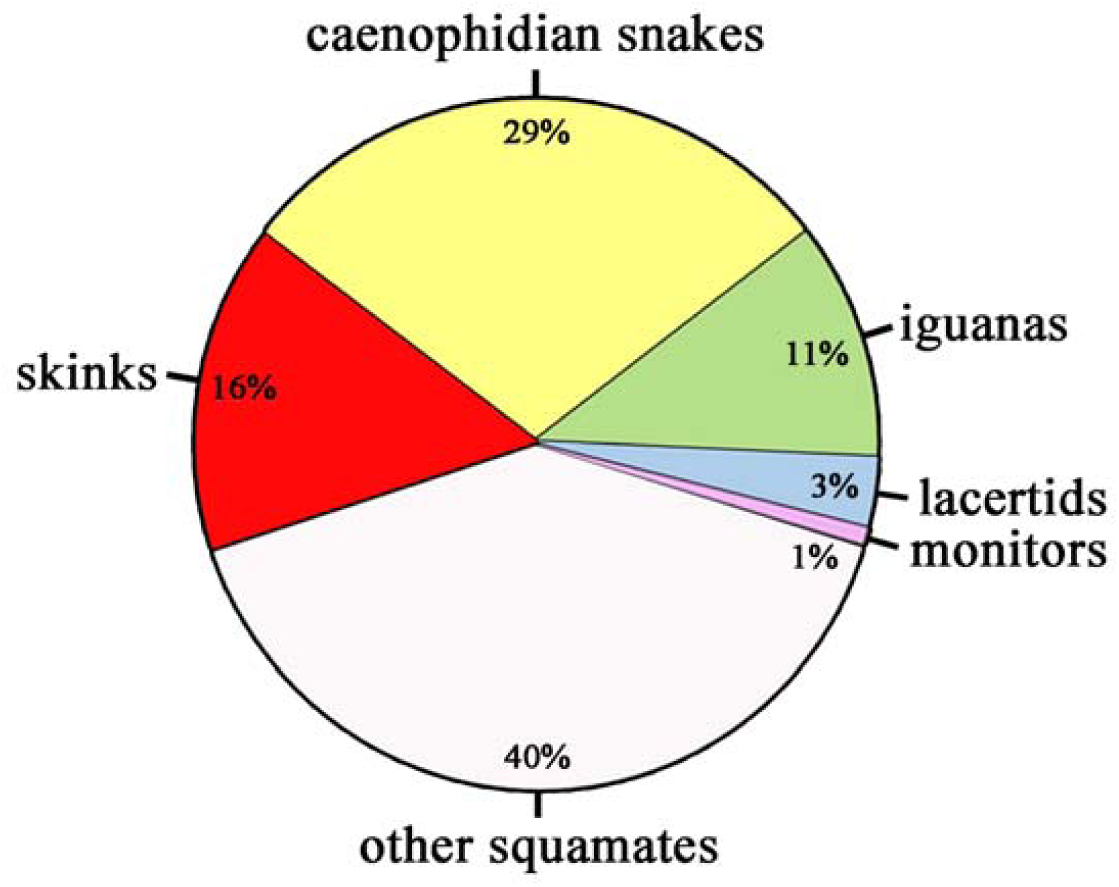
Although sex determination was traditionally assumed to be highly unstable in squamate reptiles, around 60% of squamates are members of five clades with sex chromosomes conserved for dozens of millions of years [current study;10,17-22]. A molecular sexing method based on qPCR was developed for each of these clades, allowing the accurate identification of the sex in approximately 6.000 species of squamates.

### Molecular sexing of skinks

The technique based on quantitative PCR for testing the homology of sex chromosomes across skinks can be used as a widely applicable method for molecular sexing [17]. Such methods were not available up to now in skinks, although it is very much needed. Molecular sexing is essential for many developmental and ecological studies requiring determination of the sex of embryos, which were previously hampered in this important group by the lack of a molecular sexing methods. Moreover, many skinks are extremely difficult to be sexed based on external morphology even as adults. 56 skink species are critically endangered and 77 species are endangered worldwide [80]. Our technique for molecular sexing could be crucial for the success of conservation projects. Furthermore, molecular sexing is a valuable tool for studies in developmental biology. This molecular sexing method based on qPCR is now available for potentially 60% of squamate species (figure 4).

### Conclusions

The observed homology of sex chromosomes in skinks was not expected based on recent models of sex chromosome evolution which postulated that poorly differentiated sex chromosomes are young and unstable [81], i.e. prone to turnovers or even transitions to ESD where sex chromosomes are not present. Skinks can be considered as another reptile group [figure 4] with stability of sex chromosomes. Skinks provide further support that the phylogenetic border between unstable and stable sex chromosomes does not lay between ectotherms and endotherms, but perhaps between anamniotes and amniotes [20]. The reasons for this difference remain still unknown but might be related to differences in offspring numbers generally associated with the level of genetic polymorphism [20]. Poorly differentiated sex chromosomes in highly diversified skinks also do not provide much support for the hypothesis that differentiated sex chromosomes are important contributors to radiation. The age of the poorly differentiated XX/XY sex chromosomes in skinks is similar to the differentiated ZZ/ZW sex chromosomes in birds and advanced snakes [10,23], which opposes the traditional theoretical predictions that the Y chromosome should degenerate faster than the W chromosome. The traditional predictions are also not supported by the empirical data in other squamate lineages. A higher degree of differentiation in lineages with female heterogamety in comparison to male heterogamety was observed within snakes [82], chameleons [36,37], and between closely related teiid and lacertid lizards (20,83). On the other hand, XX/XY sex chromosomes are highly differentiated in viviparous mammals and iguanas [with the exception of basilisks 76,84]; however, sex chromosomes in these two groups are among the oldest sex chromosomes uncovered to date in amniotes and have thus potentially had a longer time to differentiate [85,86]. Moreover, these two lineages are the only amniote lineages with a chromosome-wide dosage compensation mechanism [87–89], which might influence the rate of differentiation of sex chromosomes. In any case, the models based on local (in the sense of genomic region) and lineage-specific sexual differences in recombination deserve further attention [14].

## Acknowledgment

We would like to express our gratitude to Jana Thomayerová for technical assistance and Amy Wilson for the linguistic improvement of the manuscript. Blood or tissue samples for this study were kindly provided by Daniel Frynta, Jiří Šmíd and Jan Suchánek. Photos of skinks for figure 2 were kindly provided by Anna Bauerová, Arthur Georges, Lukáš Kubička, Tomáš Peš, Jiří Šmíd and Jana Thomayerová.

## Data access

The raw Illumina reads from mRNA-seq of all studied individuals are deposited into the NCBI BioProject database with ID PRJNA660179.

## Author contributions

AK: cytogenetic and molecular work; LK: statistics; MR: bioinformatics; MR, LK: conceived the project. All authors contributed to the final form of the manuscript and held responsible for its content.

## Figures

**Table S1:** List of reptiles used in the current study.

**Table S2:** Male to female ratios for DNA-seq read coverage in *S. scincus*. Homology of these genes to the genomes of the chicken *Gallus gallus* and the common wall lizard *Podarcis muralis* is given.

**Table S3:** List of primers and qPCR measurements of relative gene dosage in 13 species of skinks and their close outgroups.

## References

1. Johnson Pokorná M, Kratochvíl L. 2014 What was the ancestral sex-determining mechanism in amniote vertebrates? Biol. Rev. 91, 1–12. (doi: 10.1111/brv.12156)

2. Abbott JK, Nordén AK, Hansson B. 2017 Sex chromosome evolution: historical insights and future perspectives. Proc. R. Soc. B 284, 20162806. (doi: 10.1098/rspb.2016.2806)

3. Bull JJ, Charnov EL. 1985 On irreversible evolution. Evolution 39, 1149–1155. (doi: 10.2307/2408742)

4. Rinn JL, Snyder M. 2005 Sexual dimorphism in mammalian gene expression. Trends Genet. 21, 298–305. (doi: 10.1016/j.tig.2005.03.005)

5. Norris DO, Lopez KH. 2011 Hormones and Reproduction of Vertebrates. San Diego, CA: Elsevier Academic Press Inc.

6. Viets BE, Ewert MA, Talent LG, Nelson CE. 1994 Sex-determining mechanisms in squamate reptiles. J. Exp. Zool. 270, 45–56. (doi: 10.1002/jez.1402700106)

7. Bergero R, Gardner J, Bader B, Yong L, Charlesworth D. 2019 Exaggerated heterochiasmy in a fish with sex-linked male coloration polymorphisms. Proc. Natl. Acad. Sci. U.S.A. 116, 6924–6931.

8. Pipoly I, Bókony V, Kirkpatrick M, Donald PF, Székely T, Liker A. 2015 The genetic sex-determination system predicts adult sex ratios in tetrapods. Nature 527, 91–94. (doi: 10.1038/nature15380)

9. Xirocostas ZA, Everingham SE, Moles AT. 2020 The sex with the reduced sex chromosome dies earlier: a comparison across the tree of life. Biol. Lett. 16, 20190867.

10. Rovatsos M, Vukić J, Lymberakis P, Kratochvíl L. 2015 Evolutionary stability of sex chromosomes in snakes. Proc. R. Soc. B 282, 20151992. (doi: 10.1098/rspb.2015.1992)

11. Bachtrog D, Kirkpatrick M, Mank JE, McDaniel SF, Pires JC, Rice W, Valenzuela N. 2011 Are all sex chromosomes created equal? Trends Genet. 27, 350–357. (doi: 10.1016/j.tig.2011.05.005)

12. Mank JE. 2012 Small but mighty: the evolutionary dynamics of W and Y sex chromosomes. Chromosome Res. 20, 21–33. (doi: 10.1007/s10577-011-9251-2)

13. Wilson Sayres MA. 2018 Genetic diversity on the sex chromosomes. Genome Biol. Evol. 10, 1064–1078. (doi: 10.1093/gbe/evy039)

14. Sardell JM, Kirkpatrick M. 2020 Sex differences in the recombination landscape. Am. Nat. 195, 361–379. (doi: 10.1086/704943)

15. Perrin N. 2009 Sex reversal: a fountain of youth for sex chromosomes? Evolution 63, 3043–3049. (doi: 10.1111/j.1558-5646.2009.00837.x)

16. Pennell MW, Mank JE, Peichel CL. 2018 Transitions in sex determination and sex chromosomes across vertebrate species. Mol. Ecol. 27, 3950–3963. (doi: 10.1111/mec.14540)

17. Rovatsos M, Kratochvíl L. 2017 Molecular sexing applicable in 4000 species of lizards and snakes? From dream to real possibility. Methods Ecol. Evol. 8, 902–906.

18. Rovatsos M, Rehák I, Velenský P, Kratochvíl L. 2019 Shared ancient sex chromosomes in varanids, beaded lizards, and alligator lizards. Mol. Biol. Evol. 36, 1113–1120. (doi: 10.1093/molbev/msz024)

19. Rovatsos M, Praschag P, Fritz U, Kratochvíl L. 2017 Stable Cretaceous sex chromosomes enable molecular sexing in softshell turtles (Testudines: Trionychidae). Sci. Rep. 7. (doi: 10.1038/srep42150)

20. Rovatsos M, Vukić J, Altmanová M, Johnson Pokorná M, Moravec J, Kratochvíl L. 2016 Conservation of sex chromosomes in lacertid lizards. Mol. Ecol. 25, 3120–3126. (doi: 10.1111/mec.13635)

21. Rovatsos M, Pokorná M, Altmanová M, Kratochvíl L. 2014 Cretaceous park of sex determination: sex chromosomes are conserved across iguanas. Biol. Lett. 10, 20131093. (doi: 10.1098/rsbl.2013.1093)

22. Rovatsos M, Farkačová K, Altmanová M, Johnson Pokorná M, Kratochvíl L. 2019 The rise and fall of differentiated sex chromosomes in geckos. Mol. Ecol. 28, 3042–3052. (doi: 10.1111/mec.15126)

23. Zhou Q, Zhang J, Bachtrog D, An N, Huang Q, Jarvis ED, Gilbert MTP, Zhang G. 2014 Complex evolutionary trajectories of sex chromosomes across bird taxa. Science 346, 1246338. (doi: 10.1126/science.1246338)

24. Graves JAM. 2016 Did sex chromosome turnover promote divergence of the major mammal groups? BioEssays 38, 734–743. (doi: 10.1002/bies.201600019)

25. Gamble T, Coryell J, Ezaz T, Lynch J, Scantlebury DP, Zarkower D. 2015 Restriction site-associated DNA sequencing (RAD-seq) reveals an extraordinary number of transitions among gecko sex-determining systems. Mol. Biol. Evol. 32, 1296–1309. (doi: 10.1093/molbev/msv023)

26. Uetz P, Hošek J (eds). 2014 The reptile database. https://www.reptile-database.org (accessed 21 March 2020).

27. Vitt LJ, Caldwell JP. 2014 Herpetology: an introductory biology of amphibians and reptiles. Amsterdam; Boston: Academic Press.

28. Olmo E, Signorino G. 2013 Chromorep: a reptiles chromosomes database https://chromorep.univpm.it (accessed 21 March 2020)

29. Hutchinson MN, Donnellan SC. 1992 Taxonomy and genetic variation in the Australian lizards of the genus *Pseudemoia* (Scincidae: Lygosominae). J. Nat. Hist. 26, 215–264. (doi: 10.1080/00222939200770091)

30. Hardy GS. 1979 The karyotypes of two scincid lizards, and their bearing on relationships in genus *Leiolopisma* and its relatives (Scincidae: Lygosominae). N. Z. J. Zool. 6, 609–612. (doi: 10.1080/03014223.1979.10428403)

31. Wright JW. 1973 Evolution of the X1×2Y sex chromosome mechanism in the scincid lizard *Scincella laterale* (Say). Chromosoma 43, 101–108.

32. Donnellan SC. 1985 The evolution of sex chromosomes in scincid lizards. PhD Dissertation. Macquarie University, Sydney.

33. Shine R, Elphick MJ, Donnellan S. 2002 Co-occurrence of multiple, supposedly incompatible modes of sex determination in a lizard population. Ecol. Lett. 5, 486–489. (doi: 10.1046/j.1461-0248.2002.00351.x)

34. Wapstra E, Olsson M, Shine R, Edwards A, Swain R, Joss JMP. 2004 Maternal basking behaviour determines offspring sex in a viviparous reptile. Proc. R. Soc. B 271, S230–S232. (doi: 10.1098/rsbl.2003.0152)

35. Robert KA, Thompson MB. 2001 Viviparous lizard selects sex of embryos. Nature 412, 698–699. (doi: 10.1038/35089135)

36. Rovatsos M, Johnson Pokorná M, Altmanová M, Kratochvíl L. 2015 Female heterogamety in Madagascar chameleons (Squamata: Chamaeleonidae: *Furcifer*): differentiation of sex and neo-sex chromosomes. Sci. Rep. 5, 13196. (doi: 10.1038/srep13196)

37. Nielsen SV, Banks JL, Diaz RE Jr, Trainor PA, Gamble T. 2018 Dynamic sex chromosomes in Old World chameleons (Squamata: Chamaeleonidae). J. Evol. Biol. 31, 484–490. (doi: 10.1111/jeb.13242)

38. Iannucci A et al. 2019 Conserved sex chromosomes and karyotype evolution in monitor lizards (Varanidae). Heredity 123, 215–227. (doi: 10.1038/s41437-018-0179-6)

39. Rovatsos M, Vukić J, Mrugała A, Suwala G, Lymberakis P, Kratochvíl L. 2019 Little evidence for switches to environmental sex determination and turnover of sex chromosomes in lacertid lizards. Sci. Rep. 9, 7832. (doi: 10.1038/s41598-019-44192-5)

40. Caputo V, Odierna G, Aprea G. 1994 A chromosomal study of *Eumeces* and *Scincus*, primitive members of the Scincidae (Reptilia, Squamata). Boll. Zool. 61, 155–162. (doi: 10.1080/11250009409355876)

41. Pyron R, Burbrink FT, Wiens JJ. 2013 A phylogeny and revised classification of Squamata, including 4161 species of lizards and snakes. BMC Evol. Biol. 13, 93. (doi: 10.1186/1471-2148-13-93)

42. Mazzoleni S, Augstenová B, Clemente L, Auer M, Fritz U, Praschag P, Protiva T, Velenský P, Kratochvíl L, Rovatsos M. 2019 Turtles of the genera *Geoemyda* and *Pangshura* (Testudines: Geoemydidae) lack differentiated sex chromosomes: the end of a 40-year error cascade for Pangshura. PeerJ 7, e6241. (doi: 10.7717/peerj.6241)

43. Sumner AT. 1972 A simple technique for demonstrating centromeric heterochromatin. Exp. Cell Res. 75, 304–306. (doi: 10.1016/0014-4827(72)90558-7)

44. Endow SA. 1982 Polytenization of the ribosomal genes on the X and Y chromosomes of *Drosophila melanogaster*. Genetics 100, 375–385.

45. Bolger AM, Lohse M, Usadel B. 2014 Trimmomatic: a flexible trimmer for Illumina sequence data. Bioinformatics 30, 2114–2120. (doi: 10.1093/bioinformatics/btu170)

46. Andrews S. 2010 FASTQC. A quality control tool for high throughput sequence data.

47. Ewels P, Magnusson M, Lundin S, Käller M. 2016 MultiQC: summarize analysis results for multiple tools and samples in a single report. Bioinformatics 32, 3047–3048. (doi: 10.1093/bioinformatics/btw354)

48. Vicoso B, Emerson JJ, Zektser Y, Mahajan S, Bachtrog D. 2013 Comparative sex chromosome genomics in snakes: differentiation, evolutionary strata, and lack of global dosage compensation. PLOS Biol. 11, e1001643. (doi: 10.1371/journal.pbio.1001643)

49. Picard MAL, Cosseau C, Ferré S, Quack T, Grevelding CG, Couté Y, Vicoso B. 2018 Evolution of gene dosage on the Z-chromosome of schistosome parasites. eLife 7, e35684. (doi: 10.7554/elife.35684)

50. Andrade P et al. 2019 Regulatory changes in pterin and carotenoid genes underlie balanced color polymorphisms in the wall lizard. Proc. Natl. Acad. Sci. U.S.A. 116, 5633–5642. (doi: 10.1073/pnas.1820320116)

51. Ye J, Coulouris G, Zaretskaya I, Cutcutache I, Rozen S, Madden T. 2012 Primer-BLAST: A tool to design target-specific primers for polymerase chain reaction. BMC Bioinform. 13, 134. (doi: 10.1186/1471-2105-13-134)

52. Rovatsos M, Altmanová M, Pokorná MJ, Kratochvíl L. 2014 Novel X-linked genes revealed by quantitative polymerase chain reaction in the green anole, *Anolis carolinensis*. G3 4, 2107–2113. (doi: 10.1534/g3.114.014084)

53. Carré G-A, Siggers P, Xipolita M, Brindle P, Lutz B, Wells S, Greenfield A. 2017 Loss of p300 and CBP disrupts histone acetylation at the mouse *Sry* promoter and causes XY gonadal sex reversal. Hum. Mol. Genet. 27, 190–198. (doi: 10.1093/hmg/ddx398)

54. Falah N, Posey JE, Thorson W, Benke P, Tekin M, Tarshish B, Lupski JR, Harel T. 2017 22q11.2q13 duplication including *SOX10* causes sex-reversal and peripheral demyelinating neuropathy, central dysmyelinating leukodystrophy, Waardenburg syndrome, and Hirschsprung disease. Am. J. Med. Genet. 173, 1066–1070. (doi: 10.1002/ajmg.a.38109)

55. Polanco JC, Wilhelm D, Davidson T-L, Knight D, Koopman P. 2009 *Sox10* gain-of-function causes XX sex reversal in mice: implications for human 22q-linked disorders of sex development. Hum. Mol. Genet. 19, 506–516. (doi: 10.1093/hmg/ddp520)

56. Kuzmin A, Jarvi K, Lo K, Spencer L, Chow GYC, Macleod G, Wang Q, Varmuza S. 2009 Identification of potentially damaging amino acid substitutions leading to human male infertility. Biol. Reprod. 81, 319–326. (doi: 10.1095/biolreprod.109.076000)

57. Liška F, Chylíková B, Janků M, Šeda O, Vernerová Z, Pravenec M, Křen V. 2016 Splicing mutation in *Sbf1* causes nonsyndromic male infertility in the rat. Reproduction 152, 215–223. (doi: 10.1530/rep-16-0042)

58. Balla T. 2013 Phosphoinositides: Tiny lipids with giant impact on cell regulation. Physiol. Rev. 93, 1019–1137. (doi: 10.1152/physrev.00028.2012)

59. Firestein R, Nagy PL, Daly M, Huie P, Conti M, Cleary ML. 2002 Male infertility, impaired spermatogenesis, and azoospermia in mice deficient for the pseudophosphatase Sbf1. J. Clin. Invest. 109, 1165–1172. (doi: 10.1172/jci0212589)

60. Marshall Graves JA, Peichel CL. 2010 Are homologies in vertebrate sex determination due to shared ancestry or to limited options? Genome Biol. 11, 205. (doi: 10.1186/gb-2010-11-4-205)

61. O’Meally D, Ezaz T, Georges A, Sarre SD, Marshall Graves JA. 2012 Are some chromosomes particularly good at sex? Insights from amniotes. Chromosome Res. 20, 7–19. (doi: 10.1007/s10577-011-9266-8)

62. Sigeman H, Ponnikas S, Chauhan P, Dierickx E, Brooke M de L, Hansson B. 2019 Repeated sex chromosome evolution in vertebrates supported by expanded avian sex chromosomes. Proc. R. Soc. B 286, 20192051. (doi: 10.1098/rspb.2019.2051)

63. Gamble T, Castoe TA, Nielsen SV, Banks JL, Card DC, Schield DR, Schuett GW, Booth W. 2017 The discovery of XY sex chromosomes in a boa and python. Curr. Biol. 27, 2148–2153.e4. (doi: 10.1016/j.cub.2017.06.010)

64. Vicoso B, Emerson JJ, Zektser Y, Mahajan S, Bachtrog D. 2013 Comparative sex chromosome genomics in snakes: differentiation, evolutionary strata, and lack of global dosage compensation. PLOS Biol. 11, e1001643. (doi: 10.1371/journal.pbio.1001643)

65. Nielsen SV, Pinto BJ, Guzmán-Méndez IA, Gamble T. 2020 First report of sex chromosomes in night lizards (Scincoidea: Xantusiidae). J. Hered. 111, 307–313. (doi: 10.1093/jhered/esaa007)

66. Rovatsos M, Gamble T, Nielsen SV, Georges A, Ezaz T, Kratochvíl L. 2020 Do male and female heterogamety really differ in expression regulation? Lack of global dosage balance in pygopodid geckos. Phil. Trans. R. Soc., in press.

67. Cornejo-Páramo P et al. 2020 Viviparous reptile regarded to have temperature-dependent sex determination has old XY chromosomes. Genome Biol. Evol. 12, 924–930. (doi: 10.1093/gbe/evaa104)

68. Giovannotti M, Caputo V, O’Brien PCM, Lovell FL, Trifonov V, Nisi Cerioni P, Olmo E, Ferguson-Smith MA, Rens W. 2009 Skinks (Reptilia: Scincidae) Have highly conserved karyotypes as revealed by chromosome painting. Cytogenet. Genome Res. 127, 224–231. (doi: 10.1159/000295002)

69. Donnellan SC. 1991 Chromosomes of Australian lygosomine skinks (Lacertilia: Scincidae). Genetica 83, 207–222. (doi: 10.1007/bf00126227)

70. Zheng Y, Wiens JJ. 2016 Combining phylogenomic and supermatrix approaches, and a time-calibrated phylogeny for squamate reptiles (lizards and snakes) based on 52 genes and 4162 species. Mol. Phylogenetics Evol. 94, 537–547. (doi: 10.1016/j.ympev.2015.10.009)

71. Kumar S, Stecher G, Suleski M, Hedges SB. 2017 TimeTree: a resource for timelines, timetrees, and divergence times. Mol. Biol. Evol. 34, 1812–1819. (doi: 10.1093/molbev/msx116)

72. Radder RS, Quinn AE, Georges A, Sarre SD, Shine R. 2007 Genetic evidence for cooccurrence of chromosomal and thermal sex-determining systems in a lizard. Biol. Lett. 4, 176–178. (doi: 10.1098/rsbl.2007.0583)

73. Hill PL, Burridge CP, Ezaz T, Wapstra E. 2018 Conservation of sex-Linked markers among conspecific populations of a viviparous skink, *Niveoscincus ocellatus*, exhibiting genetic and temperature-dependent sex determination. Genome Biol. Evol. 10, 1079–1087. (doi: 10.1093/gbe/evy042)

74. Sutou S, Mitsui Y, Tsuchiya K. 2001 Sex determination without the Y chromosome in two Japanese rodents *Tokudaia osimensis osimensis* and *Tokudaia osimensis* spp. Mamm. Genome 12, 17–21. (doi: 10.1007/s003350010228)

75. Acosta A, Suárez-Varón G, Rodríguez-Miranda LA, Lira-Noriega A, Aguilar-Gómez D, Gutiérrez-Mariscal M, Hernández-Gallegos O, Méndez-de-la-Cruz F, Cortez D. 2019 Corytophanids replaced the pleurodont XY system with a new pair of XY chromosomes. Genome Biol. Evol. 11, 2666–2677. (doi: 10.1093/gbe/evz196)

76. Nielsen SV, Banks JL, Diaz RE Jr, Trainor PA, Gamble T. 2018 Dynamic sex chromosomes in Old World chameleons (Squamata: Chamaeleonidae). J. Evol. Biol. 31, 484–490. (doi: 10.1111/jeb.13242)

77. Rovatsos M, Johnson Pokorná M, Altmanová M, Kratochvíl L. 2016 Mixed-up sex chromosomes: identification of sex chromosomes in the X_1_X_1_X_2_X_2_/X_1_X_2_Y system of the legless lizards of the genus *Lialis* (Squamata: Gekkota: Pygopodidae). Cytogenet. Genome Res. 149, 282–289. (doi: 10.1159/000450734)

78. Gamble T, Coryell J, Ezaz T, Lynch J, Scantlebury DP, Zarkower D. 2015 Restriction site-associated DNA sequencing (RAD-seq) reveals an extraordinary number of transitions among gecko sex-determining systems. Mol. Biol. Evol. 32, 1296–1309. (doi: 10.1093/molbev/msv023)

79. Bókony V, Milne G, Pipoly I, Székely T, Liker A. 2019 Sex ratios and bimaturism differ between temperature-dependent and genetic sex-determination systems in reptiles. BMC Evol. Biol. 19, 57. (doi: 10.1186/s12862-019-1386-3)

80. IUCN 2020. The IUCN Red List of Threatened Species. Version 2020-1. https://www.iucnredlist.org. (accessed on 19 March 2020).

81. Perrin N. 2009 Sex reversal: a fountain of youth for sex chromosomes? Evolution 63, 3043–3049. (doi: 10.1111/j.1558-5646.2009.00837.x)

82. Augstenová B, Johnson Pokorná M, Altmanová M, Frynta D, Rovatsos M, Kratochvíl L. 2018 ZW, XY, and yet ZW: Sex chromosome evolution in snakes even more complicated. Evolution 72, 1701–1707. (doi: 10.1111/evo.13543)

83. Carvalho NDM, Arias FJ, da Silva FA, Schneider CH, Gross MC. 2015 Cytogenetic analyses of five amazon lizard species of the subfamilies Teiinae and Tupinambinae and review of karyotyped diversity the family Teiidae. Comp. Cytogenet. 9, 625–644. (doi: 10.3897/compcytogen.v9i4.5371)

84. Nielsen SV, Guzmán-Méndez IA, Gamble T, Blumer M, Pinto BJ, Kratochvíl L, Rovatsos M. 2019 Escaping the evolutionary trap? Sex chromosome turnover in basilisks and related lizards (Corytophanidae: Squamata). Biol. Lett. 15, 20190498. (doi: 10.1098/rsbl.2019.0498)

85. Cortez D, Marin R, Toledo-Flores D, Froidevaux L, Liechti A, Waters PD, Grützner F, Kaessmann H. 2014 Origins and functional evolution of Y chromosomes across mammals. Nature 508, 488–493. (doi: 10.1038/nature13151)

86. Altmanová M, Rovatsos M, Johnson Pokorná M, Veselý M, Wagner F, Kratochvíl L. 2018 All iguana families with the exception of basilisks share sex chromosomes. Zoology 126, 98–102. (doi: 10.1016/j.zool.2017.11.007)

87. Rupp SM, Webster TH, Olney KC, Hutchins ED, Kusumi K, Wilson Sayres MA. 2016 Evolution of dosage compensation in *Anolis carolinensis*, a reptile with XX/XY chromosomal sex determination. Genome Biol. Evol. 9, 231–240. (doi: 10.1093/gbe/evw263)

88. Marin R et al. 2017 Convergent origination of a *Drosophila*-like dosage compensation mechanism in a reptile lineage. Genome Res. 27, 1974–1987. (doi: 10.1101/gr.223727.117)

89. Graves JAM. 2015 Evolution of vertebrate sex chromosomes and dosage compensation. Nat. Rev. Genet. 17, 33–46. (doi: 10.1038/nrg.2015.2)

